# Importance of the Unseen: Near-Infrared Light Regulates Human Glucose Levels in the Built Environment

**DOI:** 10.64898/2026.09.06.747571

**Authors:** Khorshid Meihami, Jemima Unwin Teji, Edward Barrett, Glen Jeffery

## Abstract

Life evolved under broad spectrum sunlight (300-2500nm), but in the modern built environment under standard Light Emitting Diode (LED) illumination is restricted to around 400-650nm. Different wavelengths influence metabolism via mitochondria. 670-900nm support mitochondrial function while those in the Soret band (420-450nm) undermine mitochondrial ability. Under LED lighting, metabolism is challenged with reduced mitochondria ability, and hence, reduced demand for blood sugars that are required to produce adenosine triphosphates (ATP). As longer wavelengths enhance mitochondrial performance, they increase demand for blood sugars, reducing serum glucose. Here we change the lighting in windowless offices providing either standard LED desk lamps or those supplemented with 850nm in real-world environments. This was for two hours every morning over three days. We then assessed blood sugars with two independent methods, an oral glucose tolerance test and continuous glucose monitoring. Subjects exposed to standard LEDs had significantly increased blood sugars, while those exposed to lamps with added 850nm had significantly reduced blood sugars. These real-world data highlight the importance of light in controlling metabolism. They raise issues concerning public health in interior environments where we now spend >90% of our lives.

## Introduction

Sunlight under which life evolved has a broad spectrum from ultraviolet (UV 10-400nm) to infrared (IR 700-2500nm). Spectral elements in sunlight differentially influence metabolic activity by modulating mitochondrial function. Broad spectrum and longer wavelength exposure that penetrates deeply maintains mitochondrial function by supporting their membrane potential and promoting production of adenosine triphosphate (ATP), which fuels physiological processes [1,2]. However, shorter wavelengths on their own undermine mitochondrial function and ATP production leading to reduced physiological performance [3]. This is particularly true for wavelengths in the Soret band (420-450nm) [4,5].

Light in the built environment is dominated by shorter wavelengths than found in solar spectrum. They are generated by light emitting diodes (LEDs), which emit predominantly in the 400-650 nm range with minimal output above 700 nm [6]. This spectral restriction may reduce mitochondrial function, diminish cellular glucose demand, and predispose individuals towards impaired glycaemic regulation, a state resembling prediabetes [5,7]. In support of this, Powner and Jeffery demonstrated in controlled laboratory settings that brief exposures to 670 nm light significantly reduced blood glucose in human and animal subjects as measured by Oral Glucose Tolerance Test (OGTT), and that these effects were likely mediated via mitochondrial mechanisms [8,9]. Complementary animal studies using 670 nm and 420 nm light exposures demonstrated reciprocal effects on systemic glucose metabolism, consistent with wavelength-dependent modulation of mitochondrial function [4,10].

Despite this growing body of laboratory evidence, no study has examined whether modifying the spectral composition of everyday office lighting by adding near infrared (NIR) wavelengths to standard LED luminaires can produce measurable changes in blood glucose regulation in a real-world setting. This gap is of public health relevance as populations in the Western world now spend approximately 90–95% of their time indoors, where illumination is increasingly provided by LED lighting with spectral characteristics that differ substantially from natural daylight. However, the metabolic consequences of this environmental shift remain poorly characterised.

The application of longer wavelengths at 670nm in laboratory settings can improve glucose regulation [9]. Hence, here, we translate these findings by changing lighting in real-world windowless office environments illuminated with LEDs. We supplement standard LED desk lamps with 850 nm NIR LEDs and ask if this will significantly reduce postprandial blood glucose area under the curve (AUC) analysis compared with standard LED lighting alone, as measured by Oral Glucose Tolerance Test (OGTT) over a three-day intervention period. The 850 nm near-infrared wavelength was selected due to its ability to penetrate deeper into biological tissues while not being visually detected [1].

## Methods

### Study design

This study employed a parallel-group, randomised, double-blind, controlled design with pre- and post-intervention assessments. Participants were randomly assigned to either a standard LED lighting control group (A) or 850nm lighting intervention group (B). To maintain blinding, the control and intervention desk lamps were externally indistinguishable and provided equivalent visible illumination (approximately 425 lux; 2700 K CCT). The intervention lamp additionally emitted 850 nm NIR radiation, which is not detectable by the human visual system. Lamp labels were concealed by an independent third party before deployment, ensuring that neither participants nor researchers could identify the allocated lighting condition. The primary outcome measure was the area under the curve (AUC) for the Oral Glucose Tolerance Test (OGTT), assessed at pre and post lighting conditions exposure.

### Participants

Twenty-four healthy male participants, working in windowless offices were recruited following approval from the University College London Ethics Committee (Project ID: 0251). All research reported was performed in accordance with all relevant guidelines and regulations. The research was performed in accordance the Declaration of Helsinki. All participants provided fully informed consent. Healthy was defined as having no diagnosed medical conditions and taking no prescribed or regular medications. Female participants were excluded to minimise the potential confounding effects of cyclical hormonal fluctuations on glucose metabolism. Participants were randomly allocated to one of two groups: Group A (Control; n = 12) or Group B (850 nm NIR intervention; n = 12). Participant characteristics are presented in Table 1.

**Table 1.** Descriptive characteristics of the study sample at baseline. Values are presented as group means for the overall sample (N = 24), the Control group (n = 12), and the Intervention group (n = 12). Variables include demographic characteristics (age and body mass index [BMI]) and self-reported lifestyle factors recorded during the study period, including sleep duration on the night preceding testing, daily alcohol consumption, caffeine intake, and water consumption. BMI was calculated as weight (kg) divided by height squared (m²). Alcohol intake is reported as units per day, caffeine intake as cups per day, and water consumption as litres per day. Baseline values indicate that the Control and Intervention groups were broadly comparable across measured characteristics. These data were collected through a diary completed by participants over the course of one week.

| Variable | Overall | Control | Intervention |
| --- | --- | --- | --- |
| Age | 35.7 | 36.8 | 34.6 |
| Body Mass Index (BMI) | 25.1 | 26.1 | 24.2 |
| Hours slept the night before | 7 | 7.1 | 6.9 |
| Alcohol units per day | 0.6 | 0.6 | 0.6 |
| Caffeine cups per day | 1.8 | 1.8 | 1.8 |
| Water litre per day | 1.6 | 1.5 | 1.7 |
|  | N=24 | N=12 | N=12 |

### Sample size calculation

Sample size of 24 participants (12 per group) provides approximately 80% statistical power to detect a meaningful between-group difference of 25 mg/dL in glucose levels (Cohen’s d = 1.25; two-tailed α = 0.05) [16]. The effect size estimate of d = 1.25 was derived from the prior laboratory study by Powner and Jeffery (2024) [9], which reported a large effect of 670 nm light on glucose AUC in healthy participants. This sample size is appropriate for a proof-of-concept investigation.

### Lighting Conditions

Two identical desk lamps with E27 bulbs were used in both control and intervention group. Control group (A) was a standard E27 LED bulb; Intervention group (B) incorporated an additional 850 nm NIR LED, which is undetectable to the human eye (Deep Red, Light Power Health UK). Desk lamps were positioned on the left side of each participant by their computer screens at 50 cm diagonally from the centre of the desk and were switched on for 2 hours every morning for three consecutive days. The 2 hour exposure was selected to span the temporal window of postprandial glucose change assessed by OGTT. The three-day intervention was chosen to minimise disruption to participants normal work patterns. (Figures 1 and 2). Both lamps provided equivalent visible illumination of approximately 425 lux at a nominal correlated colour temperature (CCT) of 2700K. Radiometric measurements were taken at the work plane and at participants’ eye and head levels (Table 2).

**Figure 1.**
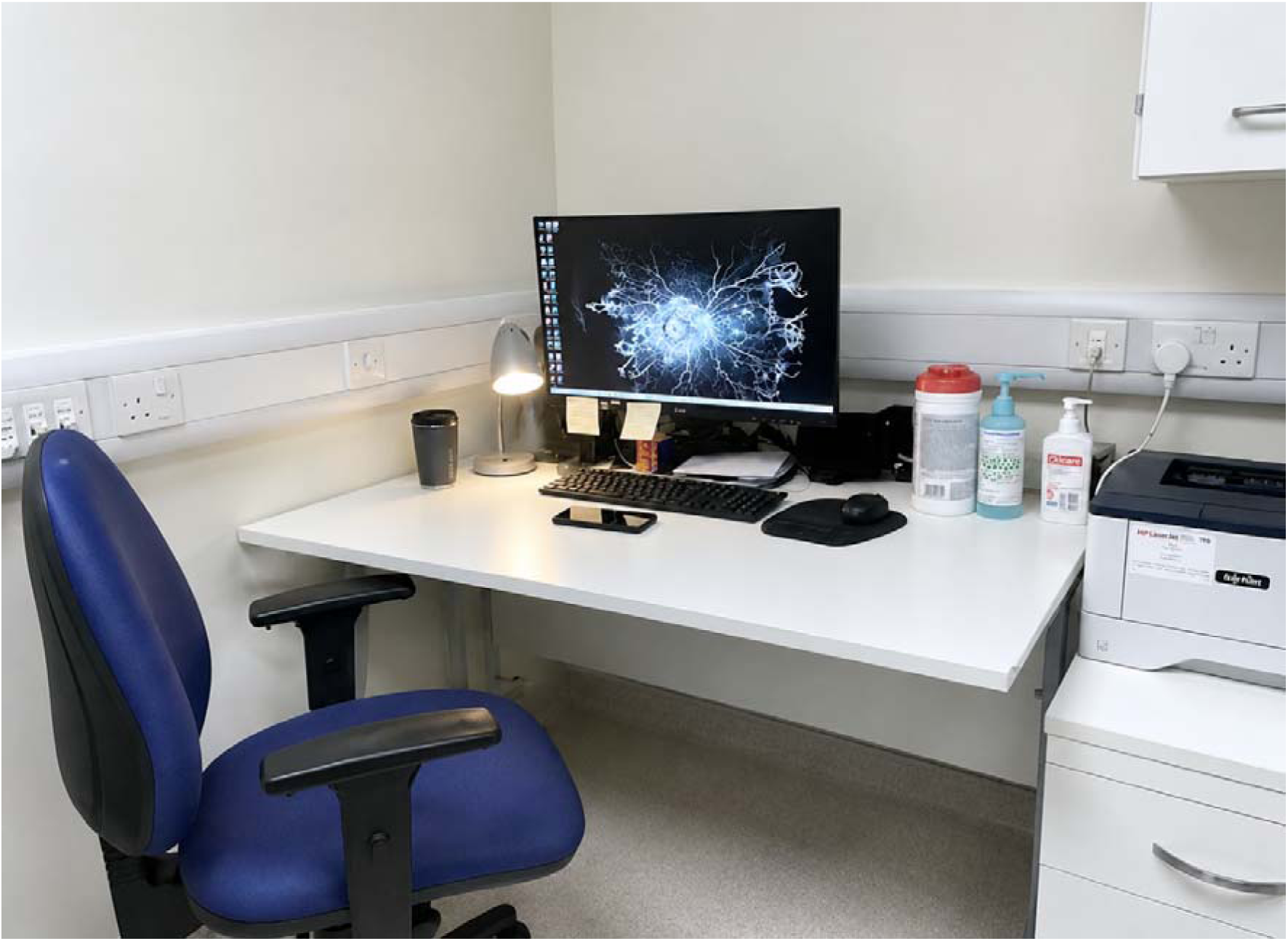
Experimental setup within the office environment. The study was conducted in a windowless office room without access to natural daylight and illuminated solely by artificial LED lighting. The desk lamp was positioned to the left of the participant’s workstation, approximately 50 cm diagonally from the centre of the desk. Participants were either in control group (A) or intervention group (B) (850 nm NIR) for 2 hours daily across three consecutive days.

**Figure 2.**
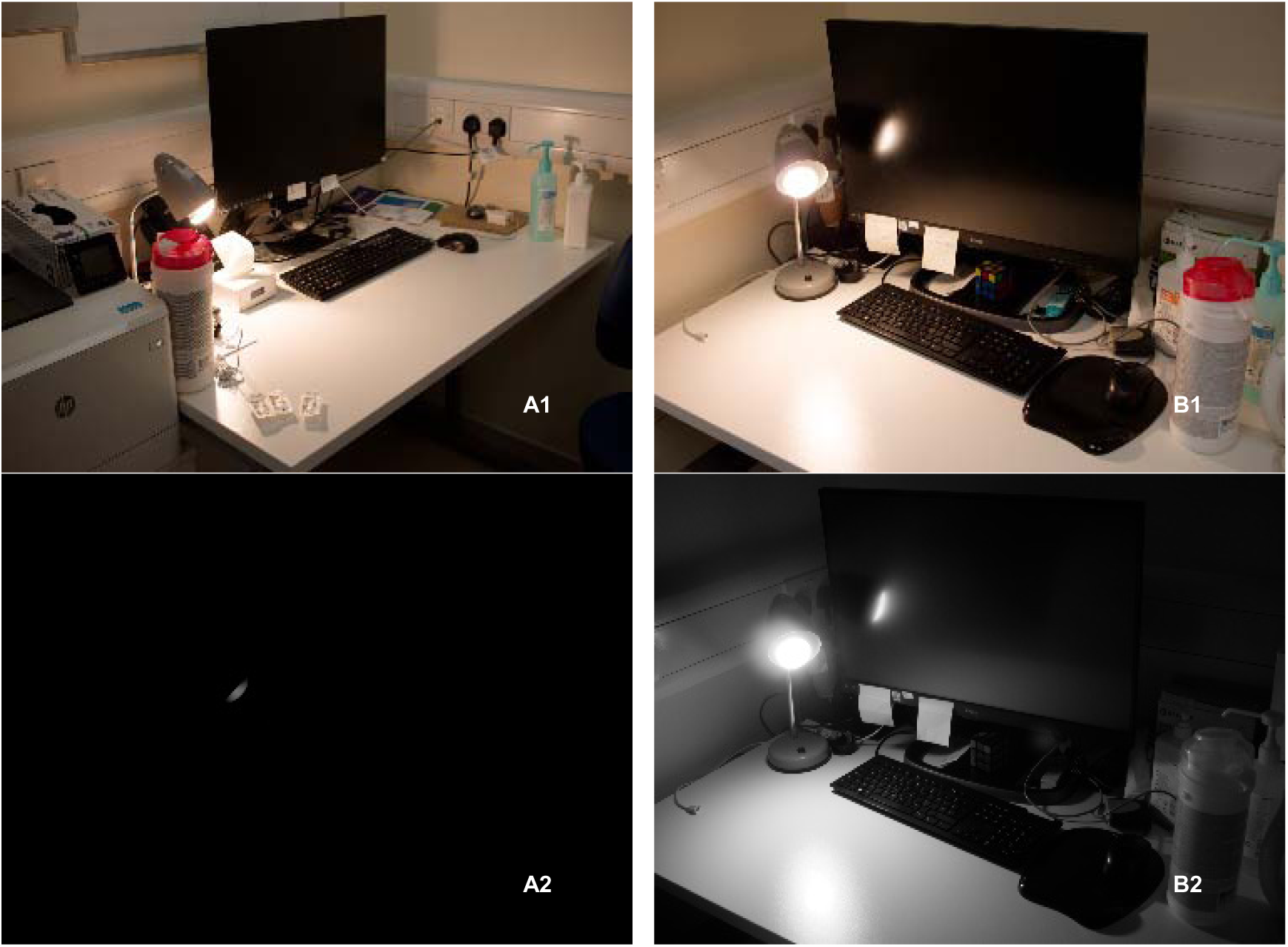
Experimental lighting conditions in the control and intervention workstations. Representative photographs of the study workstations under the two lighting conditions. (A1) Standard digital photograph of the control workstation illuminated by the visible desk lamp only, with no near-infrared (NIR) illumination. (B1) Standard digital photograph of the intervention workstation illuminated by the same visible desk lamp supplemented with an 850 nm NIR light source. Because 850 nm radiation is outside the visible spectrum, the NIR illumination is not distinguishable in conventional digital photographs. (A2) Infrared (IR) camera image of the control group with the LED lamp without 850nm switched on, confirming the absence of NIR emission. (B2) IR camera image of the intervention group lamp with 850 nm NIR source switched on, demonstrating the presence of NIR illumination. The visible lighting conditions were maintained as closely as possible between groups, with the addition of the 850 nm NIR source constituting the only experimental difference.

**Table 2.** Spectral power measurements of the lighting conditions used in the study. Lighting condition in control group (A) consisted of LED illumination without NIR, whereas lighting condition in intervention group (B) included NIR with a peak wavelength of 850 nm. These measurements were taken at desk level where the desk lamp was positioned at 50cm diagonally on the left side of the subjects. Abbreviations: CCT: Correlated Colour Temperature NIR: near infrared, LED: light emitting diode.

| Characteristics | Existing Lighting | no NIR - A | NIR Light - B |
| --- | --- | --- | --- |
| Total Irradiance (mW/cm<sup>2</sup> ) | 0.012 | 0.164 | 0.395 |
| Irradiance<750 nm (mW/cm<sup>2</sup> ) | 0.012 | 0.161 | 0.141 |
| Irradiance>750 nm (mW/cm<sup>2</sup> ) | 0 | 0.003 | 0.254 |
| Photon Flux (μmol/m<sup>2</sup>/s) | 0.52 | 5.47 | 6.54 |
| Illuminance | 50 | 425 lx | 428 lx |
| CCT | 3779 | 2700 K | 2700 K |

### Office Environments

Windowless office environments in London were specifically chosen as experimental sites because they lacked access to natural daylight. Each office was occupied by a male professional and had a mean floor area of approximately 7 m². Office environments had standard humidity and temperature regulation that were controlled centrally. The office walls were made of plasterboard and the ceiling had conventional suspended tiling. All were finished with matte white paint, providing diffuse reflective surfaces with high visible-light reflectance and minimal specular reflection. The existing lighting consisted exclusively of ceiling-mounted LED panel luminaires controlled mostly by an occupancy-sensing system. The additional desk lamps increased the total irradiance experienced by subjects by approximately more than one log unit.

### Procedure

Prior to the start of the study, a Continuous Glucose Monitor (CGM; Abbott FreeStyle Libre 2) was applied to the back of each participant’s non-dominant upper arm. On Day 1, participants attended a morning testing session and completed a standardised fasting OGTT. The subsequent three days served as a baseline monitoring period, during which continuous glucose data were collected to establish individual glucose profiles. Participants were then exposed to a three-day lighting condition using either control group (A) or intervention group (B). A second OGTT was performed on the final day of the study to assess changes in glucose regulation.

Glucose regulation was assessed using two complementary methods: continuous glucose monitoring (CGM) and oral glucose tolerance testing (OGTT). Interstitial glucose concentrations were continuously measured throughout the study using the Abbott FreeStyle Libre 2 CGM system, providing an independent assessment of glycaemic regulation alongside the OGTT. The sensor measured interstitial glucose continuously and stored glucose values at 15-minute intervals for analysis. CGM data were used to characterise glucose dynamics throughout the study, including mean daily glucose concentrations and temporal glucose profiles during both the baseline and intervention periods.

Standardised OGTTs were conducted at baseline (pre-intervention) and following the lighting intervention (post-intervention). Participants completed a 10-hour overnight fast before each test, with water permitted ad libitum. Tests were performed between 08:00 and 10:00 h on day one, before any light exposure and on the final day of the study after three days of morning light intervention, the desk lamp was switched on for 2 hours. A fasting capillary blood sample was obtained by finger-prick (0 min), after which participants consumed a standardised solution containing 75 g of glucose dissolved in 150 mL of water within 2 min. Additional capillary blood samples were collected at 15, 30, 45, 60, 75, 90, 105, and 120 min following glucose ingestion.

Throughout the study, participants continued their usual work routines and were instructed to maintain consistent lifestyle behaviour patterns. Participants were also asked to maintain their regular diet throughout the one-week study period, avoid any sudden or substantial dietary changes, and report any unusual food or beverage consumption in their daily diary. To monitor potential confounding factors, participants completed a daily diary for seven consecutive days, recording alcohol consumption, caffeine intake, water intake, sleep duration, sleep quality, perceived stress, and any deviations from their usual dietary intake (Table 1).

### Statistical Analysis

A Mann-Whitney U test was used given the small sample size (n = 12 per group) and absence of assumptions regarding normality of data distribution. All tests were two-way with a significance threshold of α = 0.05. All analyses were performed in Python 3.10.

## Results

### Primary Outcome OGTT

Analysis of the OGTT data demonstrated that supplementation with 850 nm NIR light consistently and significantly reduced postprandial blood glucose AUC, whereas exposure to standard LED lighting alone resulted in an increase in blood glucose AUC.

The Intervention group demonstrated a mean AUC change of −104.19 ± 68.58 mmol/L (SEM = 20.38; 95% CI: −150.28 to −58.10) (median = −88.88; range: −251.25 to −28.50), demonstrating a consistent reduction in postprandial glucose level following NIR light exposure. Notably, all 12 participants in the intervention group exhibited a decrease in AUC, reflecting a uniform directional response. This difference was statistically significant (p-value < 0.001).

In contrast, the Control group had a mean AUC change of +60.94 ± 81.26 mmol/L (SEM = 23.46; 95% CI: +9.31 to +112.57) (median = +54.75; range: −30.75 to +202.50), reflecting an increase in glucose over the same period. This difference was statistically significant (p-value < 0.001). (Table 3 and Figure 4)

**Figure 3.**
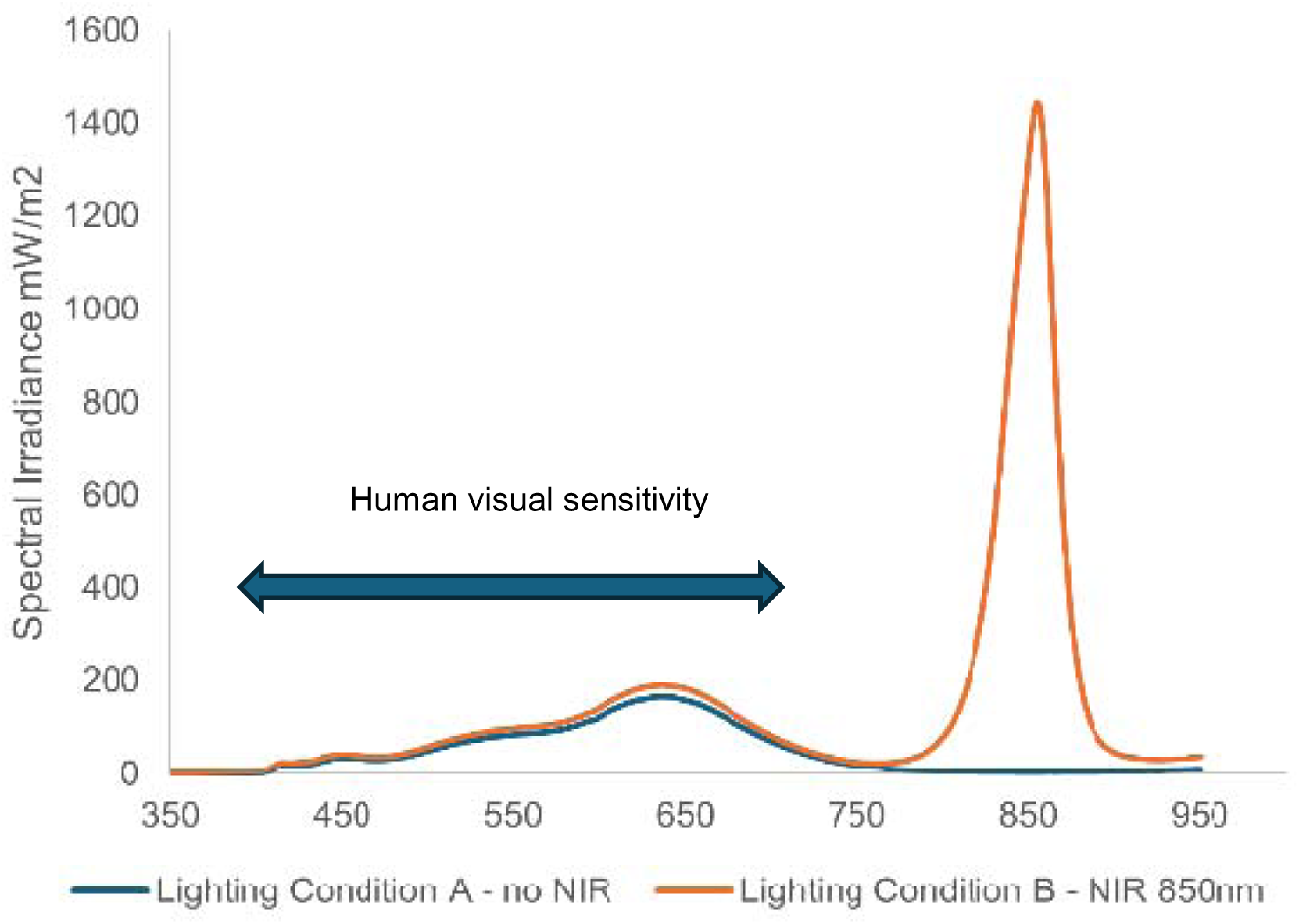
Spectral power distributions of the two lighting conditions used in the study. Lighting in Control Group (A) (blue line) consisted of standard LED illumination without NIR emission. Lighting in the intervention group (B) (orange line) included an additional NIR component with a peak wavelength of approximately 850nm. Spectral irradiance was measured at desk level at the participant workstation, with the lamp positioned approximately 50 cm diagonally to the left of the participant. The visible-light spectra of the two conditions were highly similar. Condition B exhibited a pronounced NIR peak at 850nm not present in Condition A. The blue horizontal bar represents the sensitivity of the human visual range which spans approximately 400-700nm.

**Figure 4.**
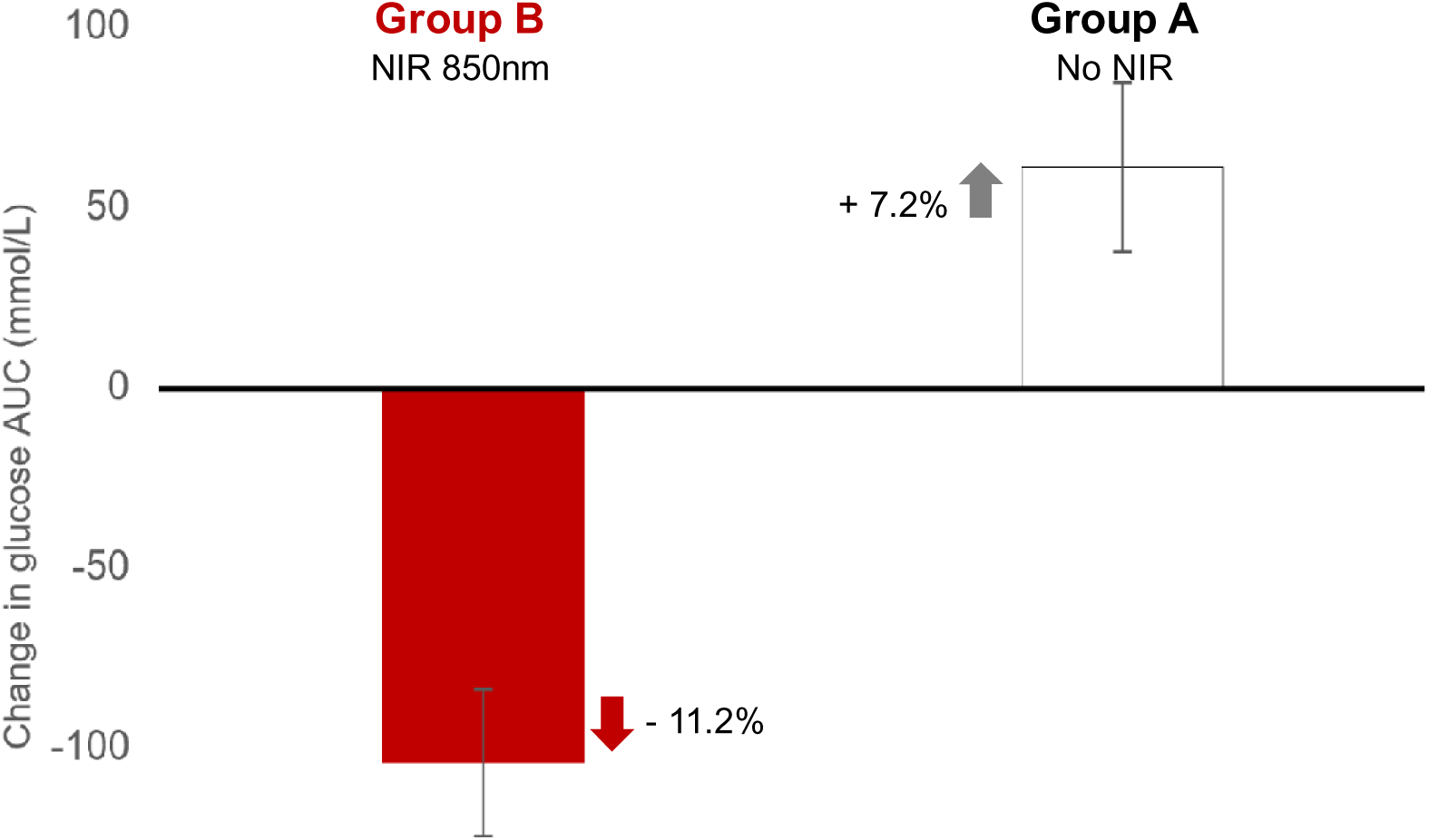
Mean change in glucose area under the curve (AUC; mmol/L) from pre-to post-intervention in the control group (A) no near infrared and the intervention group (B) with near infrared at 850nm. Error bars represent standard error of the mean (SEM). Negative values indicate a reduction in glucose AUC, whereas positive values indicate an increase. The Intervention group demonstrated a significantly greater reduction in glucose AUC compared with the Control group (Mann–Whitney U = 2.0, p < .001).

**Table 3.** Descriptive statistics and Mann–Whitney U test results for the primary outcome, change in oral glucose tolerance test (OGTT) glucose area under the curve (AUC) from pre-to post-intervention. Values are presented as mean ± standard error of the mean (SEM), with standard deviation (SD) shown in parentheses, together with median and range. AUC change was calculated as post-intervention AUC minus pre-intervention AUC. Negative values indicate a reduction in glucose exposure following the intervention (improved glucose tolerance), whereas positive values indicate an increase in glucose exposure. Between-group differences were assessed using the Mann–Whitney U test. Effect size is reported as the rank-biserial correlation (r), with larger absolute values indicating stronger group separation. The intervention group exhibited a substantial reduction in glucose AUC compared with the control group, which showed an increase in glucose AUC following the study period. OGTT: oral glucose tolerance test, AUC: area under the curve, SD: standard deviation, SEM: standard error of mean.

| Variable | Group | n | Mean ± SEM(SD) | Range | p-value |
| --- | --- | --- | --- | --- | --- |
| AUC Change<br>(Post – Pre) | Intervention | 12 | -104.19 ±<br>20.38(70.6) | -251.25 to<br>-28.50 | < 0.001 |
| AUC Change<br>(Post – Pre) | Control | 12 | +60.94 ±<br>23.46(81.3) | -30.75 to<br>+202.50 | < 0.001 |

### Secondary Outcome

Average daily glucose concentrations derived from continuous glucose monitoring (CGM) are shown in Figure 6. Participants in the intervention group exhibited a reduction in average daily glucose concentrations following exposure to 850 nm near-infrared (NIR) light during the final three days of the study. In contrast, participants in the control group demonstrated either a slight increase or minimal change in average daily glucose concentrations over the same period. The direction of change observed in the CGM data was consistent with the OGTT findings, supporting the observation that repeated 850 nm NIR exposure was associated with improved glycaemic regulation under free-living conditions. (Table 4 and Figure 6).

**Figure 5.**
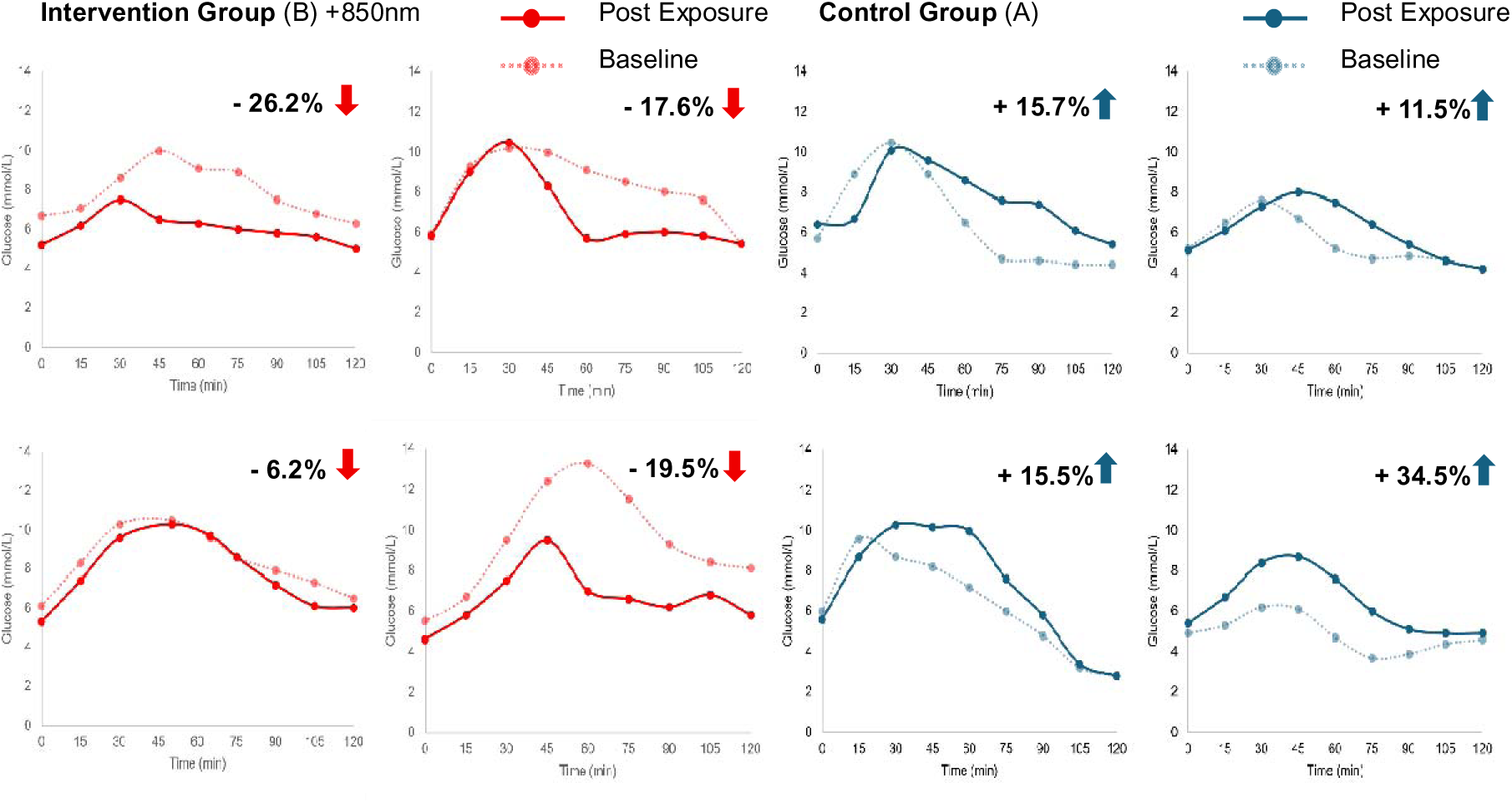
Oral Glucose Tolerance Test on Day One (baseline) Compared to Post Exposure (final day). Individual Oral Glucose Tolerance Test (OGTT) responses for eight participants before and after the near-infrared (NIR) intervention. The four panels on the left represent participants in the intervention group (B), while the four panels on the right represent participants in the control group (A). The Intervention group demonstrated a significantly greater reduction in glucose AUC compared with the Control group (Mann–Whitney U = 2.0, p < .001).

**Figure 6.**
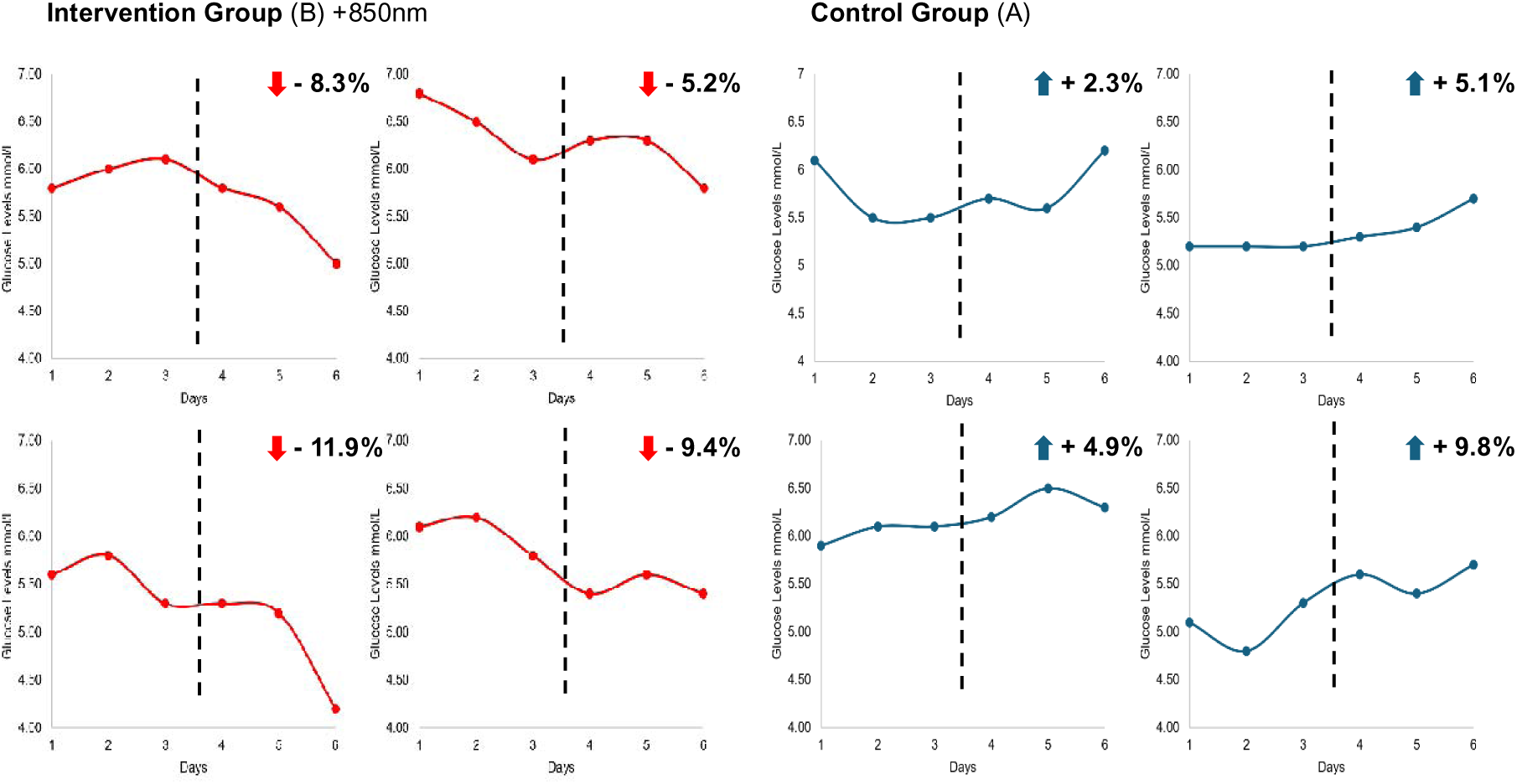
Continuous Glucose Monitoring - Average Daily Glucose over six days. Average daily glucose concentrations measured by continuous glucose monitoring (CGM) in the intervention and control groups. The intervention group (B) (left panel), exposed to 850 nm near-infrared (NIR) light during the final three days of the study period, exhibited a reduction in average daily glucose levels. In contrast, the control group (A) (right panel), showed either a slight increase or minimal change in average daily glucose levels over the same period. The vertical dashed line represents the point of intervention, with introduction of the either A or B lighting conditions.

**Table 4.** Descriptive statistics for the change in mean daily glucose concentration measured by continuous glucose monitoring (CGM) between the baseline period (Days 1–4) and the intervention period (Days 5–7). CGM data were collected at 15-minute intervals throughout the study. Values represent the difference between intervention-period and baseline-period mean daily glucose concentrations and are presented as mean change ± standard error of the mean (SEM), with standard deviation (SD) shown in parentheses. Negative values indicate a reduction in mean daily glucose concentration during the intervention period, whereas positive values indicate an increase. Data are shown for participants with complete CGM records available for analysis (n = 11 per group). Control Group (A) Intervention Group (B) CGM: continuous glucose monitor, SEM: standard error of mean, SD: standard deviation.

| Variable | Group | n | Mean Change ± SEM(SD) |
| --- | --- | --- | --- |
| Mean daily glucose | Intervention | 11 | $-0.32 \pm 0.07(0.24)$ |
| | Control | 11 | $+0.18 \pm 0.07(0.25)$ |

## Discussion

Here we reveal that standard LED lighting significantly increases blood glucose in real world office environments, while the addition of 850 nm has the opposite effect of significantly reducing blood sugars. These changes were demonstrated with two independent glucose monitoring methods, which provided consistent and complementary results.

These findings are consistent with laboratory-based human and animal studies, demonstrating that brief exposures to 670 nm light reduced blood glucose as measured by OGTT, with effects detectable at 45 minutes post light exposure [8,9]. The present study extends these findings to a longer invisible wavelength (850nm) in a real-world office environment and a multi-day exposure protocol.

The results are also consistent with other animal studies in which exposures to 670 nm reduced glucose levels while exposures to 420 nm increased them [8]. Studies in mice using 420 or 450 nm exposures which are common features of the LED spectra also showed rapid weight gain and disrupted cytokine profiles [4], consistent with the broader hypothesis that spectrally restricted LED lighting adversely affects metabolic health. While light in the Soret band is linked to these metabolic changes, it remains unclear whether the main driver for impaired glucose regulation is limited to the peak energy at 450nm or the complete absence of any infrared in LED lighting [6]. It is likely that both spectral ranges have roles to play in this key physiological function.

A second area that remains unknown is how much of the body surface needs to be exposed to obtain impact? Evidence here indicates that this can be relatively small as found by Powner and Jeffery who only illuminated an area of 800 cm^2^ region of upper back to 670nm light for 15 min at an intensity of 40mW/m−2 (28 800 J) to obtain significant systemic effect on blood glucose [9]. It is known that mitochondria act as a community across the body [11] and this community action is likely to underpin the significant reduction in blood sugars from only a relatively small regional exposure to the light. Further support for this systemic impact can be found when 850nm is shone on the back of subjects when their eyes are shielded. This resulted in subsequent improvements in vision [1]. The mechanism underpinning this systemic response may partly be found in changes in serum cytokines [12], but it is unlikely to be the only factor regulating mitochondrial communication across the body.

Lempesis et al. also showed that exposure to natural daylight for a few days can improve glycaemic control in patients with type 2 diabetes [13]. Natural daylight covers a spectral range of approximately 300–2500 nm. where long and short wavelengths have had a stable consistent relationship for billions of years, and it is likely that much of human physiology is adapted to this. Whilst consistent with the present findings, this study was limited by a small sample size, absence of spectral measurements of the environment, and the potential confounding effects of window glass filtering. The present study addresses several of these limitations through controlled spectral characterisation of the light sources and a windowless office setting that eliminates solar spectral contamination.

Full spectrum solar radiation clearly impacts on mitochondrial function, although the precise mechanism of action remains an active area of debate. It was originally thought that longer wavelengths were absorbed by cytochrome c oxidase, which is the final complex in the mitochondrial respiratory chain [14]. But this remains contentious because, longer wavelengths impact on mitochondrial respiration in *in vitro* where cytochrome c oxidase is deleted [15]. Hence, this cannot be correct.

A second proposition has been that longer wavelengths are absorbed by nano-water around mitochondria [16,17]. This is consistent with the spectral absorption characteristics of water that favours longer wavelengths. This absorption may increase the rota speed of the ATP rota pump which in turn may trigger protein synthesis that generates more complex proteins and hence more ATP [18]. This may explain the extended effect of exposures of limited duration [19].

It is likely that the mechanism resides in the action of multiple diverse biophysical pathways that do not depend on the absorption by single elements. As the fundamental mechanism relates to how electrons move through mitochondria respiratory complexes, an appreciation of Marcus theory of electron transfers may be relevant [20]. This is the current focus for debate.

This real world study reveals that LED lights supplemented with 850nm improves human metabolism. Given the large proportion of time people spend under standard LEDs, this may represent a matter of concern for public health.

## Competing interests

The authors have no competing interests

## Funding

This study received no direct funding

## Data availability

All data sets used and/or analysed during the current study are openly available at https://doi.org/10.5281/zenodo.21379180.

## Notes

https://doi.org/10.5281/zenodo.21379180.

### Competing Interest Statement

The authors have declared no competing interest.

## References

1. Jeffery, G., Hogg, C., Shinhmar, H. et al. Longer wavelengths in sunlight pass through the human body and have a systemic impact which improves vision. Scientific Reports 15, 24435 (2025). 10.1038/s41598-025-09785-3

2. Gkotsi, D., Begum, R., Salt, T., Lascaratos, G., Hogg, C., Chau, K.-Y., Schapira, A. H. V. & Jeffery, G. Recharging mitochondrial batteries in old eyes: Near infra-red increases ATP. Experimental Eye Research 122, 50–53 (2014). 10.1016/j.exer.2014.02.023

3. Kaynezhad, P., Fosbury, R., Hogg, C., Tachtsidis, I., Sivaprasad, S. & Jeffery, G. Near infrared spectroscopy reveals instability in retinal mitochondrial metabolism and haemodynamics with blue light exposure at environmental levels. Journal of Biophotonics 15, e202100283 (2022). 10.1002/jbio.202100283

4. Al-Hussaini, H., Al-Onaizi, M., Abed, B. S., Powner, M. B., Hasan, S. M. & Jeffery, G. Impact of short wavelength light exposure on body weight, mobility, anxiety like behaviour and cytokine expression. Scientific Reports 15, 5927 (2025). 10.1038/s41598-025-89081-2

5. Kam, J. H., Hogg, C., Fosbury, R., Shinhmar, H. & Jeffery, G. Mitochondria are specifically vulnerable to 420 nm light in *Drosophila* which undermines their function and is associated with reduced fly mobility. PLoS ONE 16, e0257149 (2021). 10.1371/journal.pone.0257149

6. Lei, J., Gu, X. & Liu, M. Spectral assemblage using light emitting diodes to obtain specified lighting characteristics. Applied Optics 53, 8151–8156 (2014). 10.1364/AO.53.008151

7. Linares, S. N., Beltrame, T., Galdino, G. A. M. et al. Dose response effect of photobiomodulation on hemodynamic responses and glucose levels in men with type 2 diabetes. Photonics 9, 481 (2022). 10.3390/photonics9070481

8. Powner, M. B. & Jeffery, G. Systemic glucose levels are modulated by specific wavelengths in the solar light spectrum that shift mitochondrial metabolism. PLoS ONE 17, e0276937 (2022). 10.1371/journal.pone.0276937

9. Powner, M. B. & Jeffery, G. Light stimulation of mitochondria reduces blood glucose levels. Journal of Biophotonics 17, e202300521 (2024). 10.1002/jbio.202300521

10. Almeida Rocha, M. S., Oliveira, R. S., Xavier, D. M., et al. The effects of different doses of single- and dual-wavelength whole-body photobiomodulation on metabolic parameters in obese mice. Photochemistry and Photobiology (2024). 10.1111/php.14041

11. Picard, M. & Sandi, C. The social nature of mitochondria: Implications for human health. Neuroscience & Biobehavioral Reviews 120, 595–610 (2021). 10.1016/j.neubiorev.2020.04.017

12. Shinhmar, H., Hogg, C. & Jeffery, G. Exposure to long wavelength light that improves aged mitochondrial function shifts acute cytokine expression in serum and the retina. PLoS ONE 18, e0284172 (2023). 10.1371/journal.pone.0284172

13. Lempesis, I. G. & Scheer, F. A. J. L. Illuminating the influence of natural daylight on human metabolism. Cell Metabolism 38, 7–8 (2026). 10.1016/j.cmet.2025.11.017

14. Karu, T. Primary and secondary mechanisms of action of visible to near-IR radiation on cells. J. Photochem. Photobiol. B 49, 1–17 (1999)

15. Lima, P. L. V., Pereira, C. V., Nissanka, N., Arguello, T., Gavini, G., Maranduba, C. M. da C., Diaz, F. & Moraes, C. T. Photobiomodulation enhancement of cell proliferation at 660 nm does not require cytochrome c oxidase. Journal of Photochemistry and Photobiology B: Biology 194, 71–75 (2019). 10.1016/j.jphotobiol.2019.03.015

16. Sommer, A. P. Mitochondrial cytochrome c oxidase is not the primary acceptor for near infrared light—it is mitochondrial bound water: The principles of low-level light therapy. Annals of Translational Medicine 7(Suppl. 1), S13 (2019). 10.21037/atm.2019.01.43

17. Sommer, A. P., Haddad, M. K. & Fecht, H.-J. Light effect on water viscosity: Implication for ATP biosynthesis. Scientific Reports 5, 12029 (2015). 10.1038/srep12029

18. Begum, R., Powner, M. B., Hudson, N., Hogg, C. & Jeffery, G. Treatment with 670 nm light up regulates cytochrome C oxidase expression and reduces inflammation in an age-related macular degeneration model. PLoS ONE 8, e57828 (2013). 10.1371/journal.pone.0057828

19. Shinhmar, H., Hogg, C., Neveu, M. & Jeffery, G. Weeklong improved colour contrasts sensitivity after single 670 nm exposures associated with enhanced mitochondrial function. Scientific Reports 11, 22872 (2021). 10.1038/s41598-021-02311-1

20. Zhuang, C., Qian, N. & Min, W. Quantum tunneling on water. I. General framework for microdroplet redox chemistry. The Journal of Chemical Physics 164, 224709 (2026). 10.1063/5.0326949

